# Cisplatin toxicity is mediated by direct binding to TLR4 through a mechanism that is distinct from metal allergens

**DOI:** 10.1101/2022.07.06.498982

**Authors:** Ivan K. Domingo, Jody Groenendyk, Marek Michalak, Amit P. Bhavsar

## Abstract

Cisplatin is an effective chemotherapeutic agent, yet its use is limited by several adverse drug reactions, known as cisplatin-induced toxicities (CITs). We recently demonstrated that cisplatin could elicit pro-inflammatory responses associated with CITs through Toll-like Receptor 4 (TLR4). TLR4 is best recognized for binding bacterial lipopolysaccharide (LPS) via its coreceptor, MD-2. TLR4 is also proposed to directly bind transition metals, such as nickel. Little is known about the nature of the cisplatin-TLR4 interaction. Here, we show that soluble TLR4 was capable of blocking cisplatin-induced, but not LPS-induced TLR4 activation. Cisplatin and nickel, but not LPS, were able to directly bind soluble TLR4 in a microscale thermophoresis binding assay. Interestingly, TLR4 histidine variants that abolish nickel binding, reduced, but did not eliminate, cisplatin-induced TLR4 activation. This was corroborated by binding data that showed cisplatin, but not nickel, could directly bind mouse TLR4 that lacks these histidine residues. Altogether, our findings suggest that TLR4 can directly bind cisplatin in a manner that is enhanced by, but not dependent on, histidine residues that facilitate binding to transition metals.

## INTRODUCTION

Cisplatin is the oldest and most potent of the platinum-based chemotherapeutics available (Rosenberg & VanCamp, 1970; Rosenberg *et al*, 1965). It is used to treat a variety of solid-state cancers – ranging from head-and-neck to ovarian and testicular (Nagasawa *et al*, 2021; Muzaffar *et al*, 2021; Low *et al*, 2012; Bookman, 2016; Einhorn, 2002; de Vries *et al*, 2020). Consisting of two amine ligands and two chloride ions bound to a platinum core, cisplatin mediates its anti-tumour effects by intercalating into the DNA of rapidly replicating cells. Cisplatin binding to DNA inhibits strand separation and DNA damage repair, leading to the build-up of DNA damage that eventually leads to cell death (Dasari & Tchounwou, 2014). Treatment regimens that include cisplatin can have 5-year overall survival rates of up to 90% making it integral to cancer therapy.

Despite its effectiveness, cisplatin use has been limited by the discovery of several adverse drug reactions, known as cisplatin-induced toxicities (CITs) (Tsang *et al*, 2009; El-Awady *et al*, 2011; Shahid *et al*, 2018). The development of CITs, including, but not limited to, nephrotoxicity, peripheral neurotoxicity, and ototoxicity, appears to be dependent on both the dosages administered and the age of patients. Children appear the most susceptible, particularly to cisplatin-induced hearing loss (Kamalakar *et al*, 1977; Romano *et al*, 2020; Ruggiero *et al*, 2021). Second- and third-generation platinum-based drugs, such as carboplatin and oxaliplatin, were explicitly designed with the intention of reducing those toxicities and overcoming chemotherapy resistance (Kelland, 2007). Unfortunately, the vast majority of modifications made to the underlying structure of cisplatin come at the cost of its therapeutic efficacy - leaving cisplatin the primary choice for treatment, and underscoring the need to develop mitigations for its toxicities.

Studies of the underlying mechanisms of CITs have revealed both unique features, such as anatomical location, and shared features amongst CITs. Inflammation has proven to be a critical aspect of CITs (Domingo *et al*, 2022). The inhibition of direct and indirect mediators of inflammation has proven to be effective in mitigating CITs in *in-vitro* and *in-vivo* pre-clinical studies - reducing the onset of disease as well as the overall severity (So *et al*, 2007; Babolmorad *et al*, 2021; Kim *et al*, 2011; Kaur *et al*, 2011; Rybak *et al*, 1999, 2009; Domingo *et al*, 2022). Models of CIT where pro-inflammatory signalling systems [including the Toll-like Receptor 4 (TLR4) pathway] have been removed or otherwise disabled have also exhibited considerable resistance to cisplatin toxicity (Babolmorad et al., 2021; Gao et al., 2020; Tsuruya et al., 2003; Li et al., 2019; Park et al., 2014; Ramesh & Reeves, 2003; H. S. So et al., 2008; Woller et al., 2015; B. Zhang et al., 2008; Q. Zhang et al., 2022; Y. Zhang et al., 2014; Zhou et al., 2018, 2020).

TLR4 is a membrane-bound pattern-recognition receptor (PRR) typically responsible for recognizing and mounting pro-inflammatory innate immune responses to both pathogen-associated and damage-associated molecular patterns, referred to as PAMPs & DAMPs, respectively (Kawai & Akira, 2006). Typically, TLR4 exists in complexes with co-receptors which can help dictate its specificity. The best characterized agonist of TLR4 is bacterial lipopolysaccharide (LPS), which is bound to TLR4 in conjunction with the co-receptor, MD-2. The current understanding of TLR4 activation suggests that TLR4 itself contains features that allow for the binding of various potential agonists – with co-receptors providing support by either securing target monomers for binding, or by serving as an additional binding interface to complete the TLR4 dimerization necessary for downstream signalling (Kawasaki & Kawai, 2014).

Agonists interact with TLR4 through its extracellular/ectodomain. Residues 342 to 590 of the ectodomain are highly variable between species and can confer specificity in agonist binding. An 82-amino acid ‘hypervariable’ region, found within the aforementioned ectodomain region contain unique amino acid residue combinations that in human TLR4 confers a greater affinity for hexa-acetylated LPS versus penta-acetylated LPS compared to that of murine TLR4 (Hajjar *et al*, 2002). Some of these LPS-specific amino acid residues have been characterized, such as D299 and T399, as single-nucleotide polymorphisms and found to confer LPS-specific TLR4 hypoactivation (Richard *et al*, 2021). Similarly, histidines 456 and 458 in the TLR4 ectodomain were found to mediate the interaction of TLR4 with Group 9 and Group 10 transition metals, leading to contact allergen hypersensitivities (Schmidt *et al*,2010; Raghavan *et al*, 2012; Rachmawati *et al*, 2013; Oblak *et al*, 2015). It is unclear what role these histidine residues play in mediating cisplatin-induced toxicities.

Given that platinum, the core of cisplatin, is a Group 10 transition metal as well, we hypothesized that TLR4 may be helping mediate toxicities like CITs by binding cisplatin (and platinum agents) directly and triggering hypersensitivity responses. We recently showed that genetic inhibition of Tlr4 reduced cisplatin toxicity *in vitro* and protected hair cells from cisplatin-induced death in zebrafish (Babolmorad *et al*, 2021). We further showed that TLR4 activation by cisplatin was not dependent on MD-2 and could be suppressed by TLR4 inhibitors. In this new study, we provide evidence that CITs are in part mediated by direct interactions between cisplatin and TLR4. We show that this interaction is aided, but not dependent on, key metal-binding residues in TLR4.

## RESULTS AND DISCUSSION

### Soluble recombinant TLR4 can reduce cisplatin-induced TLR4 activation *in vitro*

Platinum (II) ions, platinum (IV) ions, and cisplatin have all been shown to elicit TLR4-dependent pro-inflammatory cytokine secretion (Babolmorad *et al*, 2021). While these interactions appeared to be independent of MD-2, their molecular nature remains to be elucidated. Metal allergens, such as nickel, can elicit similar inflammatory signalling via direct activation of TLR4 (Raghavan *et al*, 2012; Schmidt *et al*, 2010; Oblak *et al*, 2015; Rachmawati *et al*, 2013). Notably, this can be blocked by soluble TLR4. Accordingly, we sought to determine whether soluble forms of TLR4 could block cisplatin activation of TLR4. HEK293 cells stably expressing human TLR4 (hTLR4), MD-2 and CD14 (HEK293-hTLR4), were pre-treated with soluble hTLR4 or mouse TLR4 (mTLR4), and then treated with either LPS, nickel or cisplatin. As expected, LPS activation of TLR4 could not be blocked by soluble hTLR4 or mTLR4, since MD-2 plays a critical role in LPS binding (Fig. 1A). Only soluble hTLR4, and not soluble mTLR4, reduced pro-inflammatory response to nickel. This was expected since mTLR4 lacks critical histidine residues found in hTLR4 that are necessary for binding Group 10 metal allergens (Schmidt *et al*, 2010). By contrast, both soluble mTLR4 and hTLR4 could inhibit TLR4 activation by cisplatin in this assay, albeit to differing extents. mTLR4 provided a 30% reduction in TLR4-dependent inflammatory cytokine secretion (Fig. 1A), while hTLR4 provided a 50% reduction (Fig. 1B). Moreover, we found that soluble hTLR4 blocking of cisplatin agonist activity was saturable (Fig. 1C).

**FIGURE 1.**
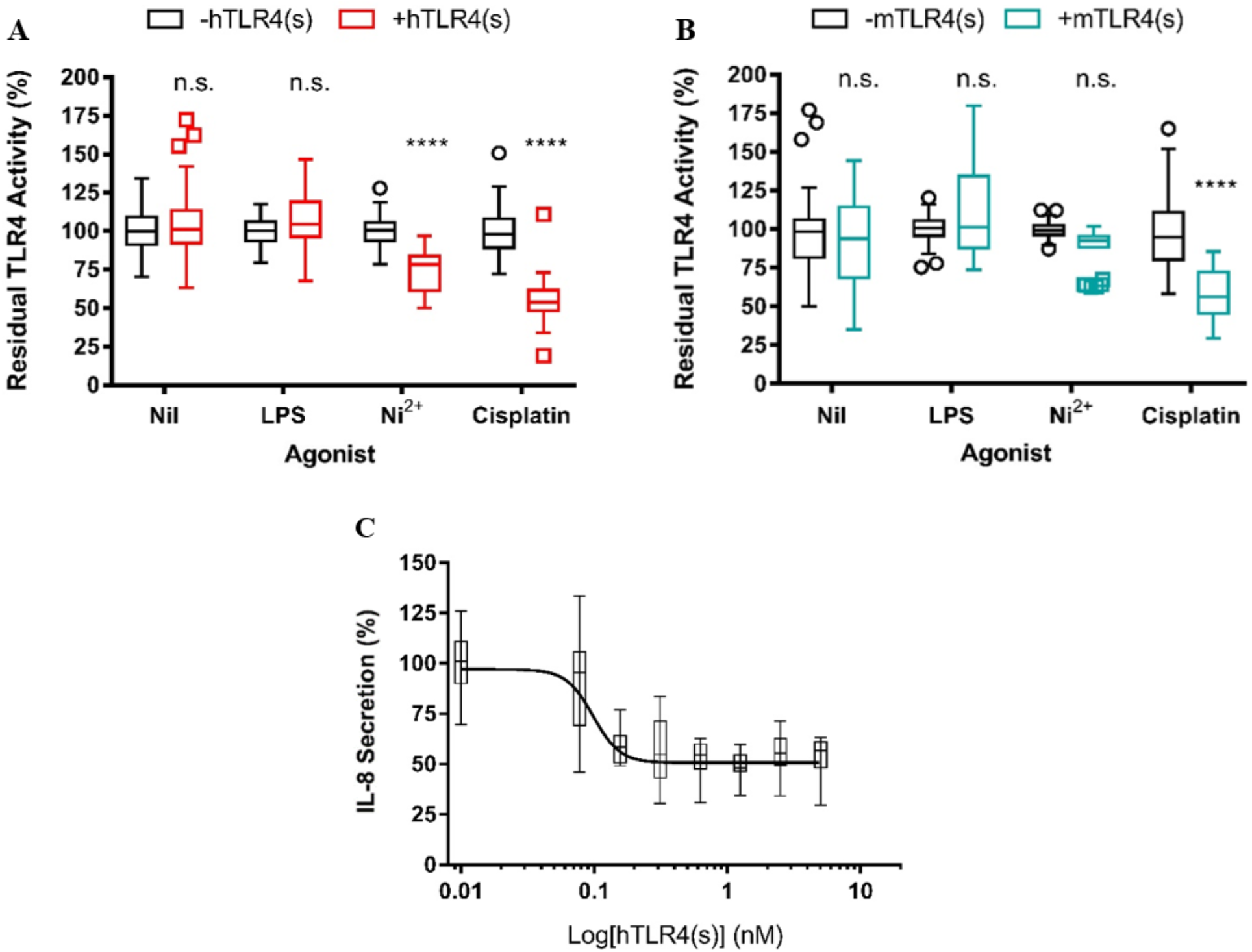
Soluble recombinant TLR4, TLR4(s), can inhibit TLR4-activated pro-inflammatory IL-8-secretion to distinct agonists in HEK293 hTLR4 cells. (A) IL-8 secretion of HEK293-hTLR4 cells pre-treated with 0.1nM soluble recombinant human TLR4 and subsequently treated with either 1 ng/mL LPS, 200 μM Ni^2+^, or 25 μM cisplatin. Data are shown as the percentage of TLR4 activity in the absence of soluble hTLR4. Nil representative of cells treated without agonists (*n* = 4 independent biological replicates for all conditions). (B) IL-8 secretion of HEK293-hTLR4 cells pre-treated with 0.1 nM soluble recombinant mouse TLR4 and subsequently treated with either 1 ng/mL LPS, 200 μM Ni^2+^, or 25 μM cisplatin. Data are shown as the percentage of TLR4 activity in the absence of soluble mTLR4 (*n* = 4 independent biological replicates for LPS & Ni^2+^ conditions, *n* = 8 independent biological replicates for Nil & Cisplatin conditions). Nil representative of cells treated without agonists. (C) IL-8 secretion of HEK293-hTLR4 cells pre-treated with different concentrations of soluble recombinant hTLR4 after treatment with 25 μM cisplatin as a percentage of IL-8 secretion without the pre-treatment condition (*n* = 4 independent biological replicates). Data Information: For all panels, actual individual data from each experiment are plotted as box (25^th^ and 75^th^ percentile borders; median central band) with Tukey whiskers. Statistical analyses were performed through 2-way ANOVA and the Bonferroni multiple testing correction. ****, *P* < 0.0001; ns, not significant. For panel C, non-linear best-fit-curve follows four parameters, variable slope; R^2^ = 0.59. Reported IC_50_ = 0.09813 nM.

### Metal allergens and cisplatin can directly bind hTLR4 without additional cellular components

Our experiments with soluble recombinant TLR4 indicate that TLR4 co-receptors are not required to block nickel and cisplatin-induced TLR4 signalling but they do not confirm a direct interaction between these ligands and TLR4. To rule out the possibility that soluble TLR4 was binding (and blocking) endogenous TLR4, or DAMPs released in response to cellular toxicity, we attempted to detect direct binding events using recombinant protein via microscale thermophoresis (MST).

To validate this system, soluble recombinant forms of hTLR4 were exposed to increasing concentrations of LPS without MD-2 based on estimated TLR4:MD-2 complex binding affinities. No detectable binding was observed between hTLR4 and LPS in this assay (Fig. 2A). By contrast, nickel was identified as a TLR4-dependent metal contact allergen, so we reasoned it could be a positive control for our MST studies, though we are not aware of any methods that have shown direct binding of nickel to TLR4. In line with previous reports, hTLR4 readily bound nickel at micromolar concentrations confirming that our methodology could detect direct TLR4 binding events (Fig. 2B).

**FIGURE 2.**
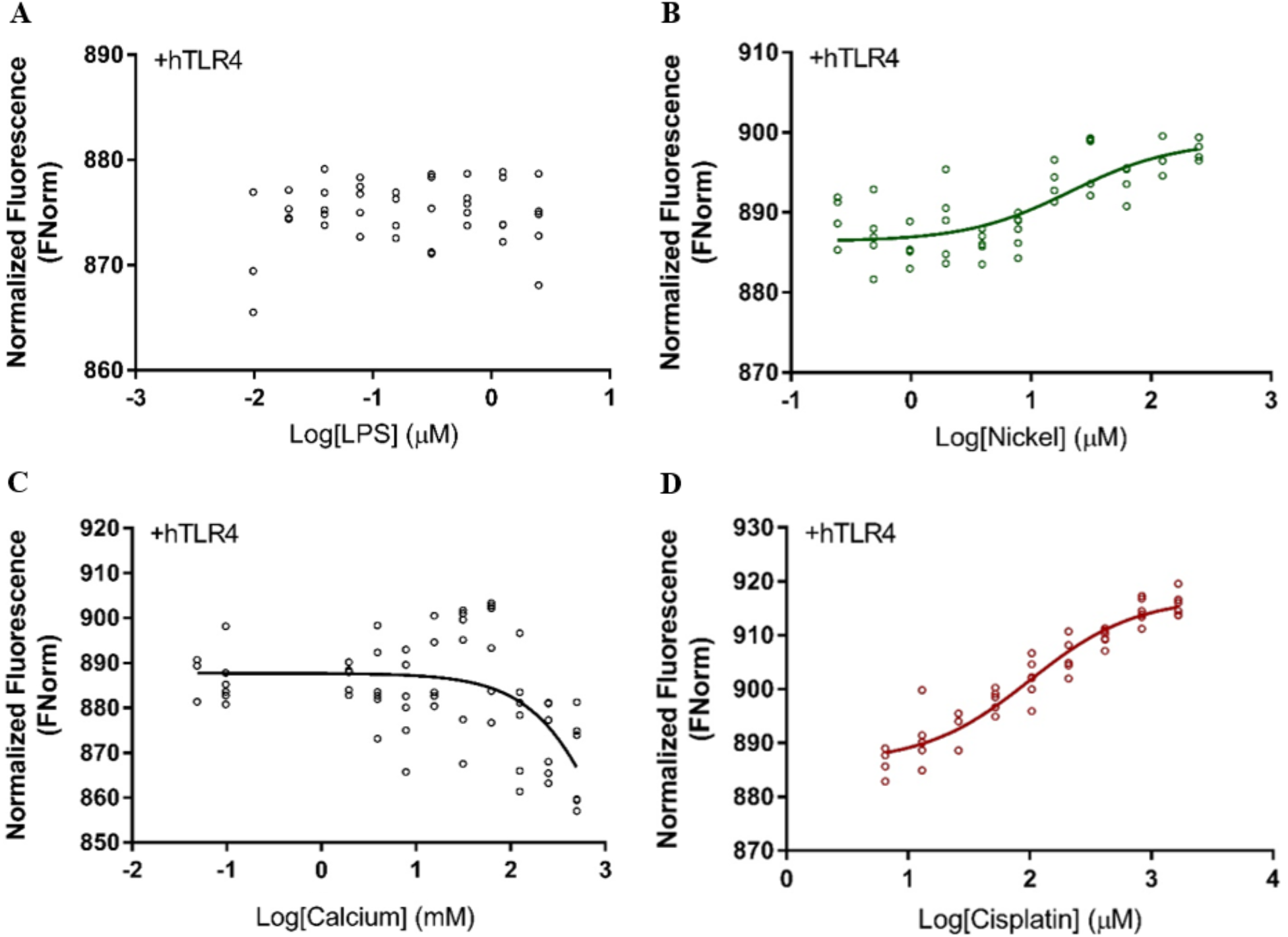
Human TLR4 can directly bind nickel and cisplatin. (A-D) Microscale thermophoresis analysis showing normalized fluorescence of hTLR4 plotted against the indicated concentrations of LPS (*n* = 5 independent replicates), Ni^2+^ (*n* = 3-6 independent replicates), Ca^2+^ (*n* = 6 independent replicates) or cisplatin (*n* = 8 independent replicates), respectively. Data Information: For all panels, data fitted to non-linear best-fit curves follow three parameters with a standardized slope. (A) Apparent Kd unavailable/undetectable. (B) Apparent K_d_ = 21.44 μM, R^2^ =0.613. (C) Apparent K_d_ = 1678 mM, R^2^ = 0.31. (D) Apparent K_d_ = 100.5 μM, R^2^ = 0.917.

To examine the specificity of our MST assay for detecting soluble TLR4 binding events, we tested calcium chloride as a ligand for TLR4. Calcium is a Group 2 metal and we are not aware of any reported calcium interactions with TLR4. Unlike nickel, hTLR4 displayed little, to no capacity, to bind calcium in our assay, even at high mM concentrations (Fig. 2C). By contrast, it has been previously shown that calcium binding to calmodulin (CaM) is robustly detected by MST (Wienken *et al*, 2010; Seeger *et al*,2017). We next tested cisplatin in the MST assay and found a clear signal of cisplatin binding to hTLR4 with a lower apparent affinity compared to nickel (Fig. 2D). Pt (II) and Pt (IV) solubility was incompatible with the MST buffer system and this precluded our ability to test their binding to hTLR4.

This work uncovers new molecular insights on the interaction between cisplatin and TLR4, supporting a role for direct activation of TLR4 by cisplatin. Our use of microscale thermophoresis with recombinant soluble TLR4 protein removes confounding factors normally associated with working with cell-based assays. Using MST, we have shown that TLR4 can bind nickel, as implied in the literature, and also cisplatin, without the need for any additional cellular factors, *e.g*. MD-2. While these findings indicate that cisplatin is sufficient to bind and activate TLR4, it does not rule out the involvement of other TLR4 agonists, e.g. LPS or DAMPs, that may contribute toTLR4 activation during CITs *in vivo*.

### TLR4 activation by platinum and cisplatin is enhanced by known metal-binding residues

To further explore these direct hTLR4-cisplatin interactions, we used site-directed mutagenesis to replace histidines 456 and 458 (located within the extracellular region of hTLR4 that is part of soluble hTLR4) with alanine and leucine, respectively. These histidines have been reported to be critical mediators of nickel-TLR4 binding (Schmidt *et al*, 2010; Raghavan *et al*, 2012). HA epitope-tagged hTLR4 variant constructs were transiently expressed in HEK293-Null2 (which do not express endogenous hTLR4, MD-2 or CD14) prior to treatment with LPS, nickel, platinum (II), platinum (IV) or cisplatin. Transiently expressed TLR4 levels were assessed by immunoblotting. hTLR4 WT and hTLR4 H456A-H458L levels were comparable, indicating these variants do not induce gross destabilization of hTLR4 (Fig. 3A). The replacement of metal-binding residues His456 and His458 had no impact on LPS-induced TLR4 activation (Fig. 3B) upon MD-2 co-expression, as expected. Also expected, the hTLR4 H456A-H458L variant was impaired in IL-8 secretion in response to nickel compared to wild-type hTLR4 and showed no significant change from the empty vector control (Fig. 3C). Interestingly, while mutation of metal-binding hTLR4 histidine residues also impaired IL-8 secretion induced by platinum (II), platinum (IV), and cisplatin (Fig. 3D, E, F), the effect appeared blunted compared to that associated with nickel. Notably, hTLR4 H456A-H458L retained significant activity when stimulated with cisplatin compared to the empty vector control. This was in stark contrast to our observations using nickel as an agonist, where activation of hTLR4 H456A-H458L was not significantly different from the empty vector control. Taken together, the data suggests that the direct interactions between cisplatin and hTLR4 occur, at least in part, due to the metal-binding properties intrinsic to hTLR4.

**FIGURE 3.**
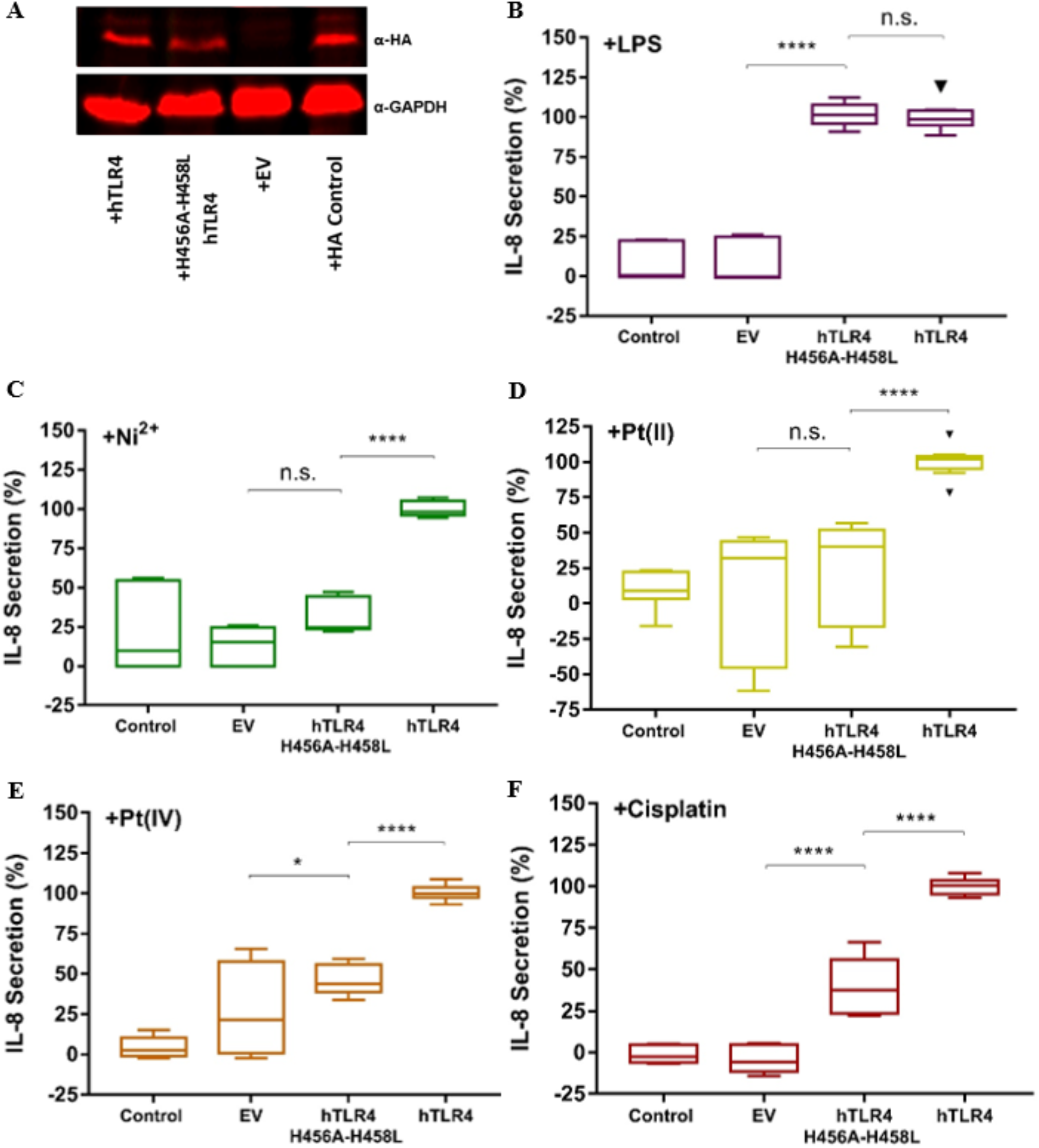
Histidine 456 & 458 mutations (H456A-H458L) inhibit TLR4 activation by platinum ions and cisplatin. (A) HEK293-Null2 cells were transfected with either empty vector (EV), hTLR4, or hTLR4 mutant constructs; immunoblotting was used to analyze relative TLR4 protein levels. (B-F) IL-8 secretion from HEK293-Null2 cells transfected with either nothing (control), empty vector (EV), hTLR4, or hTLR4 mutant constructs and subsequently treated with either 1 ng/mL LPS, 200 μM Ni^2+^, 100 μM platinum (II) [Pt(II)], 100 μM platinum (IV) [Pt(IV)], or 25 μM cisplatin displayed as a percentage of the response elicited by wild-type hTLR4 (*n* = 4 independent biological replicates for experiments performed with LPS, Ni^2+^, and Pt(II); *n* = 3, independent biological replicates for experiments involving Pt(IV) and cisplatin). Cells treated with LPS were also co-transfected with an MD-2 construct. Data Information: For panels B-F, actual individual data from each experiment are plotted as box (25^th^ and 75^th^ percentile borders; median central band) with Tukey whiskers. Statistical analyses were performed through 2-way ANOVA (for the LPS experiments) or 1-way ANOVA (for all remaining experiments) with Bonferroni multiple testing correction. *, *P* < 0.05; \*\**P* < 0.01; ***, *P* < 0.001; ****, *P* < 0.0001; n.s., non-significant.

### Cisplatin and nickel use distinct mechanisms to bind TLR4

Previous investigations have identified critical differences in how mTLR4 and hTLR4 interact with potential agonists. For example, mTLR4 does not contain conserved H456 and H458 residues. Our blocking experiments with soluble mouse TLR4 and the differential impact of His456 and His458 mutation on TLR4 activation by nickel and cisplatin hinted at differences in the ways these agonists interact with TLR4. To further explore this concept, we investigated nickel and cisplatin binding to soluble mouse TLR4 in the MST assay. Unlike our experiments with human TLR4, we were unable to detect binding of nickel to mouse TLR4 by MST (Fig. 4A). By contrast, as shown in Fig. 4B, cisplatin was able to bind to mouse TLR4 in the MST assay, albeit with a lower apparent affinity compared to human TLR4 (133 μM mTLR4 vs. 100 μM hTLR4).

**FIGURE 4.**
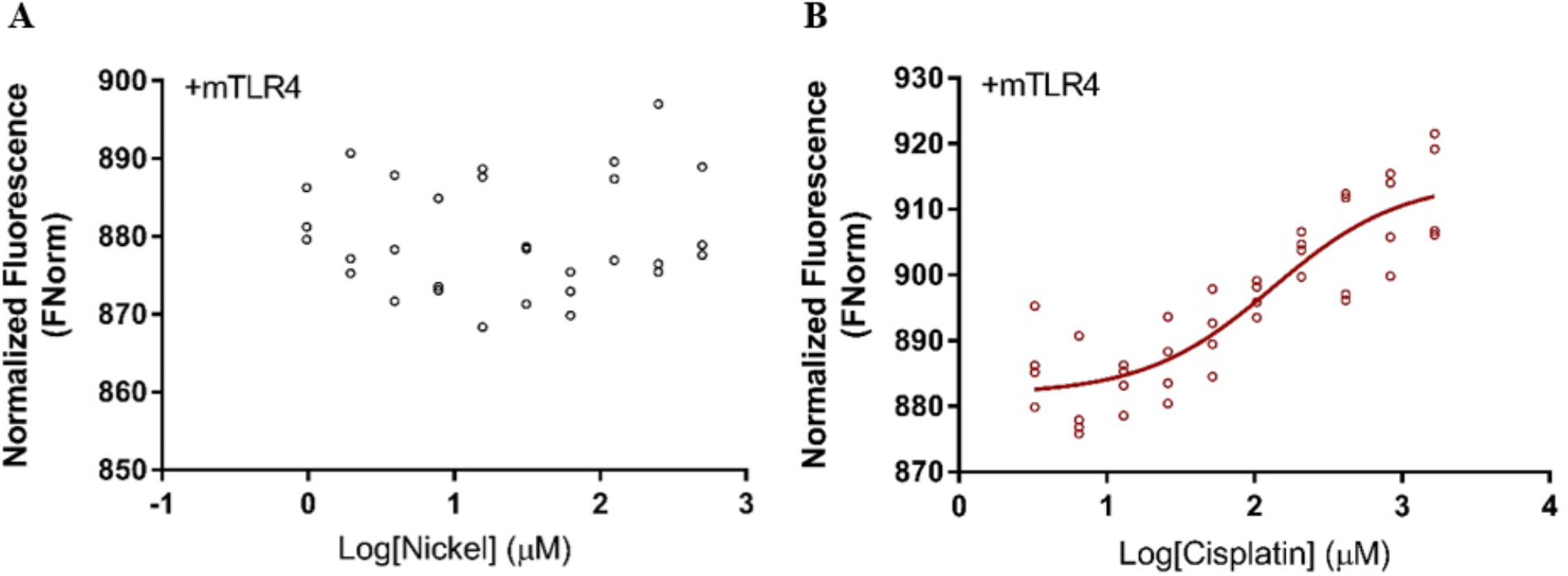
Mouse TLR4 directly binds cisplatin but not nickel. (A-B) Microscale thermophoresis analysis showing normalized fluorescence of mTLR4 plotted against the indicated concentrations of (A) Ni^2+^ (*n* = 3 independent replicates) or (B) cisplatin (*n* = 4 independent replicates), respectively. Data Information: (A) Apparent K_d_ unavailable/undetectable. (B) Apparent K_d_ = 133.6 μM; R^2^ = 0.781.

The connection between TLR4 and the development and severity of CITs has been a key area of investigation. TLR4 has been linked molecularly to CITs; TLR4 activation can induce the production of reactive oxygen and nitrogen species, apoptosis, and pro-inflammatory signalling pathways, e.g., NF-κB. Yet, the molecular details of this linkage were unclear. Initial models suggested that DNA damage generated by cisplatin could elicit the release of DAMPs that could then be subsequently detected through PRRs such as TLRs (Miller *et al*, 2010; Manohar & Leung, 2018). The relevance of TLR4-specific DAMPs to CIT development remains inconclusive, however. Cisplatin has minimal effects on TLR4-specific DAMP expression (Zhang *et al*, 2008) and in our previous work, we established that DAMP signalling can be differentiated from cisplatin-induced TLR4 activation, since TLR4 activation in response to HMGB1 depended on co-receptor, MD-2, in contrast to cisplatin (Babolmorad *et al*, 2021). Alternatively, others have posited that cisplatin-induced activity was dependent on synergies with other TLR4 ligands, such as LPS (Oh *et al*, 2011). We now add more evidence of an independent contribution to cisplatin toxicity that encompasses direct activation of TLR4 by cisplatin. Moreover, this work uncovers that this interaction is similar, but not identical, to that of metal-binding interactions resulting in known hypersensitivities. Similarly, there appears to be a critical distinction between the ability of mTLR4 and hTLR4 to bind cisplatin that requires further characterization.

In summary, our data provides evidence for direct TLR4-cisplatin binding interactions that are enhanced by, but are not dependent on, TLR4-metal-binding interactions. This is reinforced by our prior work wherein we showed that both mTLR4 and hTLR4 were mediators of cisplatin-induced ototoxicity that can be chemically targeted to mitigate cisplatin toxicity. Moreover, our findings argue for distinct mechanisms of TLR4 activation by cisplatin, nickel and LPS (Figure 5). This raises the attractive opportunity of selectively inhibiting TLR4 activation rather than ablating its function entirely. A precision treatment to reduce cisplatin-induced toxicities, while preserving TLR4 bacterial detection, would be an ideal otoprotectant to optimize the safety of cisplatin treatment.

**FIGURE 5.**
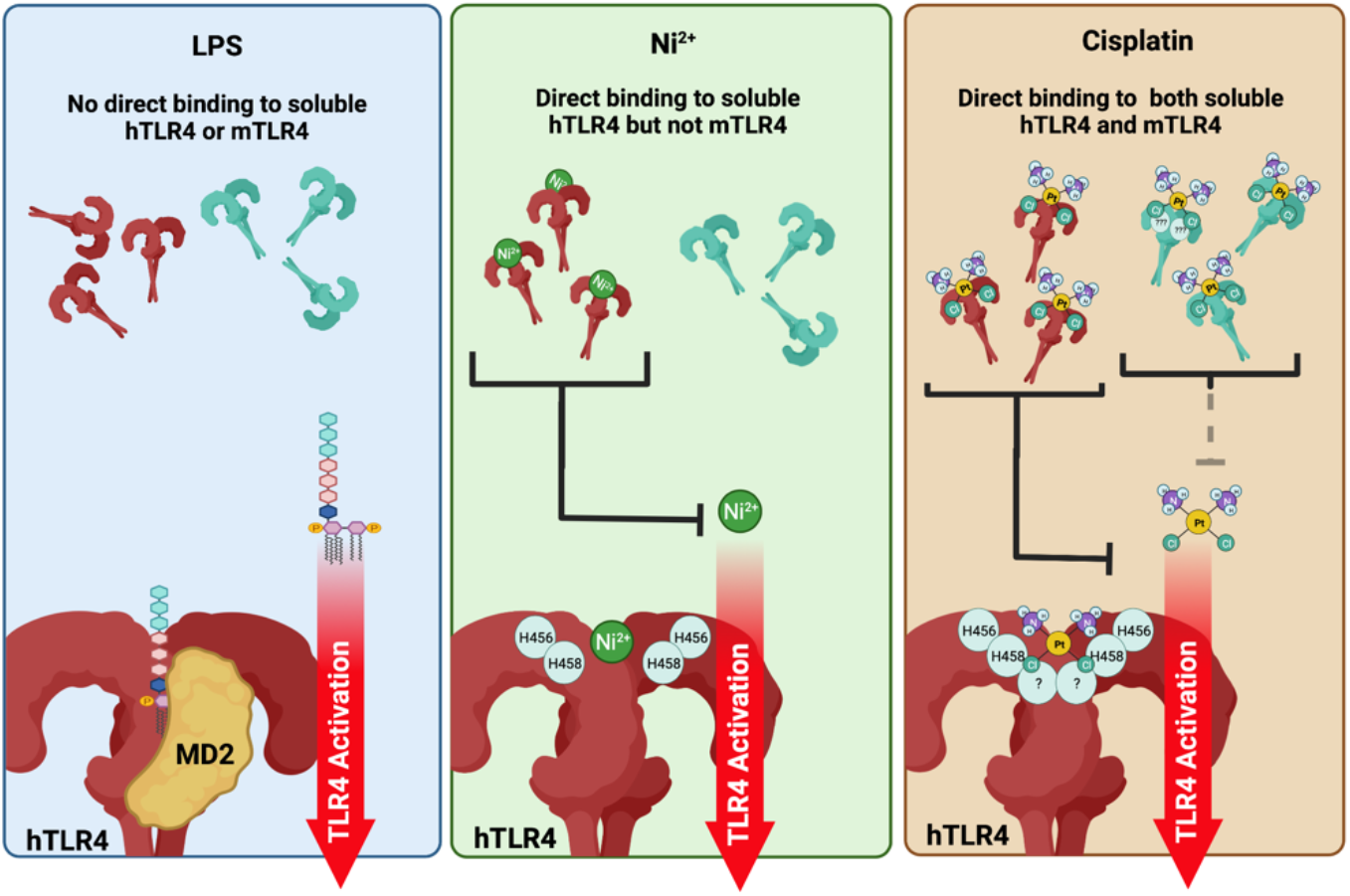
TLR4 is activated by LPS, nickel and cisplatin by distinct mechanisms. (Left panel) LPS requires MD-2 for TLR4 activation and cannot directly bind soluble TLR4 species. (Middle panel) Nickel can directly bind human TLR4 and this requires His456/His458. Soluble hTLR4 can block nickel activation of TLR4 (denoted by black line). Nickel cannot bind to mouse TLR4, which lacks these histidine residues. (Right panel) Cisplatin can directly bind both human and mouse TLR4 and this is not strictly dependent on His456/His458. Cisplatin has higher affinity for soluble hTLR4, which is more effective at blocking cisplatin activation of TLR4, compared to soluble mTLR4 (denoted by solid black vs. grey dashed lines). The additional TLR4 residues that contribute to cisplatin binding remain to be elucidated (denoted by ?).

## MATERIALS AND METHODS

### Cell Culture and Treatments

Human Embryonic Kidney HEK293-Null2 cells (Cat #hkb-null2) and HEK293-hTLR4 cells (Cat #hkb-htlr4) were obtained from Invivogen and are isogenic reporter cell lines, where HEK293-hTLR4 are stably transfected with human *TLR4* and *MD-2*. Cells were grown in DMEM supplemented with FBS (10%), penicillin-streptomycin (100μg/mL), and Normocin (100μg/mL) at 37°C and 5% CO_2_. For experiments with soluble recombinant TLR4, cells were grown in 24-well plates at a density of 5.0 × 10^4^ cells/well for 24 hrs. Either soluble mTLR4 (Biotechne R&D Systems, Cat #9149-TR-050) or hTLR4 (Biotechne R&D Systems, Cat #1478-TR-050), were added 24 hrs post-seeding for 1 hr prior to agonist treatments. Recombinant proteins were resuspended in PBS, diluted for use in culture media as described. For experiments with TLR4 histidine variants, cells were seeded in 6-well plates at 2.5 × 10^5^ cells/well. Cells were transfected 24 hrs post-seeding, where appropriate. LPS (Invitrogen, L23351), nickel chloride hexahydrate (Sigma, 654507), platinum (II) chloride (Sigma, 520632), platinum (IV) chloride (Sigma, 379840), and/or cisplatin (Teva, 02402188) treatments were made 48 hrs post-seeding, or 24hrs posttransfection. Supernatant collection, ELISAs, and cell viability analyses were performed 48 hr post-agonist treatment.

### TLR4 Histidine-456 and Histidine-458 Multi-Site Directed Mutagenesis

*TLR4* histidine residues 456 and 458 were replaced with alanine and leucine, respectively, by site-directed mutagenesis according to the manufacturer protocols ((Agilent, Cat# 200514/200515) The mutagenic primers were: 5’-TACCTTGACATTTCTGCTACTCTCACCAGAGTTGCTTTCAATGGC-3’ and 5’-GCCATTGAAAGCAACTCTGGTGAGAGTAGCAGAAATGTCAAGGTA-3’. TLR4 mutations were confirmed by Sanger sequencing using primers: 5’ - TTGGGACAACCAGCCTAAG-3’ and 5’ - GAGAGGTCCAGGAAGGTCAA-3’

### Immunoblotting

Transfected cells were collected and lysed using 400μL of Pierce RIPA Lysis and Extraction Buffer (ThermoScientific, Cat#89900), containing Pierce protease inhibitors (ThermoScientific, Cat#A32953). To lyse cells, cells were kept on ice and scraped after 15 minutes of exposure to lysis buffer. Samples were heated at 80°C for 10 min. and protein separated by SDS-PAGE prior to transfer to nitrocellulose membrane. Membranes were probed overnight with mouse anti-HA (1:2500) (Santa Cruz, 12CA5 sc-57592) or mouse anti-GAPDH (1:5000) (Invitrogen, MA5-15738) and then probed for 1 hr with goat anti-mouse secondary antibody (1:5000) (LiCor, IRDye 800CW). Probed membranes were imaged on a LiCor Odyssey and the immunoblotting procedure was performed according to their recommendations.

### Cell Transfections

HEK293-Null2 cells were transfected with either an empty vector control, human *TLR4* expression clone (kindly provided by Dr. A. Hajjar, Cleveland clinic), or *TLR4* variant expression clone. To assess the impact of histidine variants on TLR4-mediated immune responses to LPS, HEK293-Null2 cells were also co-transfected with a human MD-2 expression clone (OriGene, RC204686). JetPRIME (Polyplus, CA89129-924) reagent was used for all transfections in accordance with manufacturer specifications.

### Enzyme-Linked Immunosorbent Assay (ELISA) and Cell Viability Assays

IL-8 secretion was quantified by ELISA (88-8088, Invitrogen) as a measure of TLR4 activation in HEK-null2 and HEK-hTLR4 cells as recommended by the provider (Invivogen) Secreted cytokines were collected 48 hrs post-agonist treatment and quantified by ELISA following the manufacturer protocols. Protein secretion was normalized to cell viability to account for the differing toxicities of agonists. Cell viability was measured using MTT reagent (3-(4,5-dimethylthiazol-2-yl)-2,5-diphenyl tetrazolium bromide) (ACROS, 158990010) for the purpose of normalizing ELISA data. MTT was added to cells post-treatment at 1mg/mL and incubated for 4 hrs. The absorbance of solubilized formazan was measured at 590 nm on a SpectraMAX i3x plate reader (Molecular Devices).

### Microscale Thermophoresis (MST)

The NanoTemper Microscale Thermophoresis Monolith system was used to measure normalized fluorescence changes associated with protein-ligand binding. Soluble recombinant human or mouse TLR4 and TLR4 agonists were prepared, separately, in PBS containing 2% DMSO, and 0.5% Tween-20. Serial dilutions of agonists were mixed 1:1 with solutions of hTLR4 (100μg/mL) or mTLR4 (500μg/mL). Binding assays were performed using NanoTemper NT.LabelFree instrument (NanoTemper Technologies). All experiments were carried out at room temperature in hydrophobic capillaries with 20% light-emitting diode power (fluorescence lamp intensity) and 40% microscale thermophoresis power (infrared laser intensity). Microscale thermophoresis data were analyzed by Monolith Affinity Analysis v.2.2.6 software.

### Data Analyses and Statistical Analyses

For soluble TLR4 experiments, residual TLR4 activity was calculated by normalizing the data from soluble TLR4-treated cells to their respective control cells that lacked soluble TLR4. For investigating the dose-response to soluble hTLR4, data fitted to non-linear best fit curves following four parameters with a variable slope. 2-way ANOVA with Bonferroni multiple comparison tests were used to calculate the statistical significance of the effects of soluble TLR4 pre-treatment. To determine the percentage of IL-8 secretion retained or lost due to the mutation of histidine residues, data was normalized to the amount of IL-8 secretion triggered by the wild-type hTLR4. For experiments involving variant hTLR4 constructs, actual individual data from each experiment are plotted as boxes (25^th^ and 75^th^ percentile borders; median central band) with Tukey whiskers. Statistical analyses were performed through 2-way ANOVA (for the LPS experiments) or 1-way ANOVA (for all remaining experiments) with Bonferroni multiple testing correction For all thermophoresis experiments, data fitted to non-linear best-fit curves follow three parameters with a standardized slope. All statistical analyses were performed using Graphpad Prism 7.2.

## ACKNOWLEDGEMENTS

This work was supported by operating grants the Canadian Institutes of Health Research to APB (PJT-178327) and to MM (PS-168843). This research has also been funded by the Natural Sciences and Engineering Research Council of Canada to MM (RGPIN-2019-04908), the Li Ka Shing Institute of Virology (to APB) and the generous support of the Stollery Children’s Hospital Foundation through the Women and Children’s Health Research Institute (to APB). APB holds a Canada Research Chair in Pattern Recognition Receptor Pathophysiology and this research was undertaken, in part, thanks to funding from the Canada Research Chairs Program (101342). We are grateful to Dr. A. Hajjar for providing TLR4 expression constructs. We would additionally like to thank Dr. Harley Kurata, Dr. David Marchant, Ryley McClelland and Malcolm Forster (University of Alberta) for technical assistance with immunoblotting. Figure 5 was created with BioRender.com

## AUTHOR CONTRIBUTIONS

Conceptualization: IKD, APB, JD, MM. Investigation: IKD. Formal Analysis: IKD, APB, JG, MM. Funding Acquisition and Supervision: APB, MM. Writing - Original Draft: IKD. Editing & Revisions: All Authors.

## CONFLICTS OF INTEREST

The authors declare that they have no conflict of interest.

